# Exploring Erdr1-Mid1 Axis: Shared Risk Factors and Molecular Mechanisms in Aging and Degenerative Diseases

**DOI:** 10.1101/2025.03.07.641679

**Authors:** Yuhang Wang, Degong Pang, Ning Li

**Author notes:** **Corresponding authors**: Yuhang Wang.

## Abstract

Aging and degenerative diseases are characterized by the progressive decline in cellular, tissue, and organ function, resulting in a significant reduction in quality of life and posing major medical challenges. This highlights the urgent need to elucidate the underlying mechanisms and to develop innovative therapeutic approaches. In this study, we identify Erdr1 and Mid1 as shared risk factors for aging and multiple degenerative diseases. We propose that they contribute to disease progression by modulating oxidative stress, a well-established driver of aging and degenerative processes. We demonstrate that Erdr1 and Mid1 are both involved in oxidative stress regulation. Notably, Erdr1 undergoes alternative splicing in response to oxidative stress, resulting in reduced production of its antioxidant isoforms (Erdr1-177 and Erdr1-209), while promoting the secretion of its pro-oxidant isoform (Erdr1-145). Moreover, Erdr1-145 exacerbates oxidative damage by activating Mid1, a key inducer of oxidative stress. The Erdr1-Mid1-oxidative stress axis provides a molecular mechanistic basis for their shared role as risk factors for aging and degenerative diseases. Furthermore, we propose therapeutic strategies to mitigate cellular damage by regulating Erdr1 levels, implying a straightforward and effective approach for in situ repair of damage associated with aging and degenerative diseases.

**Highlights:**

1. Erdr1 and Mid1 are shared risk factors in aging and multiple degenerative diseases.
2. Erdr1 and Mid1 contribute to cellular damage and degeneration by modulating oxidative stress.
3. Erdr1 undergoes alternative splicing in response to oxidative stress, downregulating antioxidant isoforms (Erdr1-177 and 209) while promoting the secretion of prooxidant isoform (Erdr1-145).
4. Erdr1-145 robustly primes oxidative stress by promoting Mid1 activation, thereby highlighting the role of the Erdr1-Mid1-oxidative stress axis in cellular damage.
5. Modulating Erdr1 indicates promising strategies for in situ repair of damage associated with aging and degenerative diseases.

## Introduction

Aging and degenerative diseases are marked by a gradual decline in the structural and functional integrity of cells, tissues, and organ systems. Neurodegenerative disorders such as Alzheimer’s disease (AD)^1^, Parkinson’s disease (PD) ^2^, amyotrophic lateral sclerosis (ALS) ^3^, and Huntington’s disease (HD) ^4^ are particularly harmful due to their effects on neurons in the brain and spinal cord. Muscular degenerative conditions, such as Duchenne muscular dystrophy, cause progressive muscle weakness and atrophy ^5^, while pulmonary fibrosis (PF) is characterized by progressive scarring of lung tissue and a gradual decline in lung function ^6^. Rheumatoid arthritis (RA) is a chronic inflammatory disease characterized by progressive subchondral bone destruction^7^. Additionally, diabetes, a metabolic disorder characterized by impaired insulin production and dysregulated glucose homeostasis, is increasingly recognized as a chronic degenerative disease. This recognition stems from its progressive damage to pancreatic beta cells, often exacerbated by prolonged exposure to elevated glucose levels ^8^. Poorly controlled diabetes can cause severe complications, leading to degenerative damage in the brain, kidneys, retina, and other organs ^8,9^. Together, these degenerative diseases affect a substantial portion of the global population, highlighting the urgent need for innovative therapeutic solutions.

Oxidative stress, defined as an imbalance between the production of reactive oxygen species (ROS) and the body’s antioxidant defenses, leads to tissue and organ damage and dysfunction ^10^. It has been shown to play a critical role in the pathogenesis of aging ^11^ and various degenerative diseases, including Alzheimer’s disease (AD) ^12,13^, Parkinson’s disease (PD) ^13,14^, Huntington’s disease (HD) ^13,15^, Amyotrophic Lateral Sclerosis (ALS) ^16^, Muscular Dystrophy (MD) ^17^, Pulmonary Fibrosis (PF) ^18^, Rheumatoid Arthritis (RA) ^19,20^ and diabetes and its complications ^21,22^. Despite its well-established roles, the common intrinsic factors that regulate oxidative stress and contribute to aging and degenerative diseases remain unknown.

In this study, we identify Erdr1 and Mid1 as shared risk factors for aging and multiple degenerative diseases, proposing that they contribute to disease progression by modulating oxidative stress. We demonstrate that both Erdr1 and Mid1 play crucial roles in regulating oxidative stress. Notably, Erdr1 undergoes alternative splicing in response to oxidative stress, leading to a reduction in antioxidant isoforms (Erdr1-177 and Erdr1-209), while promoting the secretion of the pro-oxidant isoform (Erdr1-145). Moreover, Erdr1-145 exacerbates oxidative damage by activating Mid1, which serves as a key trigger for oxidative stress. The Erdr1-Mid1-oxidative stress axis provides a mechanistic explanation for their shared role as risk factors in aging and degenerative diseases. Furthermore, we propose therapeutic strategies to mitigate cellular damage by modulating Erdr1 levels, offering a straightforward and effective approach for in situ repair of damage associated with aging and degenerative diseases.

## Materials and Methods

### Cell culture

RAW 264.7 cells were cultured with RPMI-1640 medium supplemented with10% FBS, 100 IU/ml penicillin, 100 μg/ml streptomycin. 293T and HeLa cells were cultured with DMEM medium in the presence of 10% FBS, 100 IU/ml penicillin, and 100 μg/ml streptomycin.

### Induction of Cellular Oxidative Stress

To induce oxidative stress, cells were treated with different agents:

**High Glucose Treatment:** Cells were incubated in medium containing high concentrations of glucose (20–100 mM) for 24 hours to mimic a hyperglycemic environment. Control cells were cultured under normal glucose conditions (5 mM). **LPS and Nigericin Treatment:** Cells were first exposed to lipopolysaccharide (LPS, 500 ng/mL) for 3 hours, followed by a 1-hour treatment with Nigericin (10–20 µM), known to activate oxidative stress and the NLRP3 inflammasome. **H₂O₂ Treatment:** Cells were exposed to hydrogen peroxide (H₂O₂) at a final concentration of 500 µM for 4 hours. **PMA Treatment:** Cells were treated with phorbol 12-myristate 13-acetate (PMA) at a final concentration of 1 µM for 4 hours.

### Recombinant Erdr1

Recombinant Erdr1 was generated following a standard protocol. Briefly, Erdr1-209, Erdr1-177, and Erdr1-145, each with a C-terminal His-tag, were cloned into the pET28a vector. Escherichia coli (E. coli) was transformed with this plasmid and cultured in LB medium containing kanamycin. Protein expression was induced by adding isopropyl β-D-1-thiogalactopyranoside (IPTG). Erdr1-209, Erdr1-177, and Erdr1-145 were purified using a nickel column, achieving a purity of >95%, as confirmed by SDS-PAGE and immunoblot analysis. Potential endotoxin contamination was removed using High-Capacity Endotoxin Removal Spin Columns (88274, ThermoFisher).

### Anti-Erdr1 polyclonal antibody

The Anti-Erdr1 peptide antibody (36–51 amino acids of Erdr1-177, N-RAPRPPRHTRHTRHTR-C) was received as a gift from the University of Utah ^23^. This antibody can recognize all three isoforms of Erdr1.

### Plasmid preparation and transduction

The Mid1 CRISPR/Cas9 gene knockout plasmid and Mid1 expression plasmid were purchased from GenScript and cloned into the pHAGE vector. Lentivirus was produced by transfecting 293T cells with the pHAGE vector (transfer plasmid), psPAX2 (packaging plasmid), and pMD2.G (envelope plasmid) following a standard transfection protocol. After virus production, the lentiviral supernatant was used to infect RAW 264.7 cells for 48 hours.

### Immunoblotting

Following experimental treatments, both cellular and supernatant proteins were isolated for immunoblot analysis. Total cell extracts were washed twice with PBS and lysed using radioimmunoprecipitation assay (RIPA) buffer, supplemented with Complete Mini Protease Inhibitor Cocktail and Phosphatase Inhibitor Cocktail (Roche). The lysates were incubated on ice for 30 minutes, with vortexing every 5 minutes. After centrifugation at 15,000 rpm for 15 minutes at 4°C, the supernatants were collected. Proteins were resolved by SDS-PAGE and transferred to a 0.45-μm PVDF membrane. Primary antibodies against Erdr1 and β-actin were used at a 1:2000 dilution. Horseradish peroxidase (HRP)-conjugated secondary antibodies were used according to the host species of the primary antibodies.

### Immunofluorescence (IF) staining

Following experimental treatments, RAW 264.7 cells were washed twice with PBS and fixed with 4% PFA in PBS for 15 minutes at room temperature to preserve cellular structures. After fixation, cells were washed three times with PBS. Permeabilization was performed by incubating the cells with 0.1% Triton X-100 in PBS for 15 minutes at room temperature, allowing antibody penetration. The cells were then washed three times with PBS and then blocked by incubating with 5% normal goat serum in PBS for 1 hour at room temperature. The cells were then incubated overnight at 4°C with an anti-mouse Mid1 primary antibody (Catalog# PA5-38524, Invitrogen) diluted in 1% bovine serum albumin (BSA) in PBS. After incubation, the cells were washed three times with PBS to remove unbound primary antibodies. Next, Alexa Fluor 594-conjugated goat anti-mouse IgG secondary antibody (1:500 dilution) was applied to the cells and incubated for 1 hour at room temperature in the dark. Afterward, the cells were washed three times with PBS. Coverslips were mounted onto glass slides using a DAPI-containing mounting medium to stain the nuclei. Fluorescence images were acquired using a Zeiss LSM710 fluorescence microscope, ensuring consistent exposure settings across all samples for accurate comparison and analysis.

### Reactive Oxygen Species (ROS) measurement

To assess general oxidative stress, we used the CM-H2DCFDA (5-(and-6)-chloromethyl-2’,7’-dichlorodihydrofluorescein diacetate, acetyl ester) probe, a cell-permeable fluorogenic dye. Cells were seeded in a 96-well plate at a density of 4 × 10^4^ cells per well and incubated at 37°C overnight. The next day, cells were subjected to experimental treatments. Following treatment, cells were washed with PBS and incubated with 10 µM CM-H2DCFDA in serum-free medium for 30 minutes at 37°C in the dark. After incubation, cells were washed twice with PBS to remove excess dye, and fluorescence was measured using a microplate reader at an excitation wavelength of 492 nm and an emission wavelength of 527 nm. Fluorescence intensity was used for quantifying oxidative stress levels.

### LDH assay

Lactate dehydrogenase (LDH) activity in cell supernatants was measured using a commercial LDH cytotoxicity assay kit. Briefly, RAW 264.7 cells were seeded in a 6-well plate and incubated overnight before the treatment described. Following the designated treatments, cell culture supernatant was collected. A 100 µL aliquot of the supernatant was combined with 100 µL of LDH assay reagent in a 96-well plate and incubated for 30 minutes at room temperature, protected from light. Absorbance was measured at 495 nm using a microplate reader. Cytotoxicity was calculated according to the manufacturer’s instructions.

### Statistics

P values for two group comparisons were determined by Student’s t-test. p value < 0.05 was considered significant. Results were visualized using box-and-whisker plots.

## Results

### 1. Erdr1 and Mid1 are shared risk factors in aging and multiple degenerative diseases

Erythroid Differentiation Regulator 1 (Erdr1) and Midline 1 (Mid1) are evolutionarily conserved genes found in both mice and humans ^24^. The Erdr1-Mid1 axis has been demonstrated to promote pro-inflammatory processes in our previous research ^24^. In this study, we found that both Erdr1 and Mid1 are dysregulated in various cells and tissues associated with aging and multiple degenerative diseases (**Table 1; Supplemental Figures 1 and 2**), as supported by evidence from the following literature.

**Table 1.** Erdr1 and Mid1 are dysregulated in aging and various degenerative diseases.

| Diseases | Model | Cell/Tissue | Erdr1 level | Mid1 level |
| --- | --- | --- | --- | --- |
| **Senescence** | Senescence-Accelerated Prone (SAMP) strains | brain | downregulated<sup>29</sup> | upregulated<sup>30</sup> |
| **Alzheimer's Disease (AD)** | AD patients | brain | — | upregulated<sup>31-33</sup> |
|  | Mice carrying humanized APOE4 (the risk factor for AD) compared to those with APOE2 | brain | downregulated<sup>34</sup> | — |
| **Parkinson's Disease (PD)** | PD patients | brain | — | upregulated<sup>35,36</sup> |
|  | Parkin knockout mice, a PD mice model | brain | downregulated<sup>37</sup> | — |
|  | WT vs MLKL mice which exhibit resistance to PD-like symptoms | brain | downregulated<sup>38</sup> | — |
|  | A9 region (more vulnerable to neurodegeneration in PD) vs A10 region (more resistant to neurodegeneration in PD) of mice brain | neurons | downregulated<sup>39</sup> | — |
|  | 6-OHDA-induced neurotoxic mouse model of PD | brain | upregulated<sup>40</sup> | — |
| **Huntington's Disease (HD)** | CAG repeat expansions | Cell | — | Mid1 enhances the translation of HTT mRNA<sup>41</sup> |
|  | Huntington-null vs Huntington wild-type ES cell | ES cells | upregulated<sup>44</sup> | — |
|  | mutant HTT ES cells vs Huntington wild-type ES cells | ES cells | upregulated<sup>44</sup> | — |
| **Amyotrophic Lateral Sclerosis (ALS)** | Mutant SOD1-harboring NSC-34 cells | neurons | Downregulated<sup>46</sup> | — |
|  | ALS patients | neurons | — | upregulated<sup>47</sup> |
|  | ALS mouse model | neurons | — | upregulated<sup>48</sup> |
| **Muscle Degenerative Diseases (MD)** | dystrophin-deficient mouse models | heart | upregulated<sup>52</sup> | — |
|  | DMD patients | muscle | — | genomic rearrangements in Mid1<sup>54</sup> |
| **Pulmonary Fibrosis (PF)** | bleomycin-induced pulmonary fibrosis | lung | downregulated<sup>57</sup> | upregulated<sup>58</sup> |
|  | radiation-induced lung fibrosis | lung | upregulated<sup>59</sup> | upregulated<sup>59</sup> |
| **Rheumatoid Arthritis (RA)** | collagen-induced arthritis (CIA) mouse model | synovial tissue | protective role<sup>63</sup> | upregulated<sup>60</sup> |
|  | antigen-induced arthritis (AIA) mouse model | Myeloid cell | downregulated<sup>61</sup> | upregulated<sup>61</sup> |
| **Diabetes and its complications** | Insulin production upon hyperglycemia | islets | upregulated<sup>64</sup> | upregulated<sup>64</sup> |
|  | db/db diabetic mouse model | kidney | downregulated<sup>65</sup> | upregulated<sup>68</sup> |
|  | NOD mice | brain | upregulated<sup>67</sup> | upregulated<sup>67</sup> |

— represent there are no available data.

#### 1.1 Erdr1 and Mid1 are dysregulated in aging

Aging is a major risk factor for many degenerative diseases, causing a decline in the function of cells, tissues, and organs ^25,26^. Erdr1 was found to be significantly dysregulated in the brain and lung tissues of aged mice compared to their younger counterparts^27,28^. Furthermore, Erdr1 was significantly downregulated in the brain of the Senescence-Accelerated Mouse Prone 8 (SAMP8) strain ^29^, a strain characterized by accelerated aging, neuroinflammation, and oxidative stress. Meanwhile, Mid1 was upregulated in the brain of SAMP10 mice, a strain associated with a reduced lifespan and cognitive decline ^30^. Notably, Mid1 expression was significantly reduced in the hippocampus of SAMP10 mice following L-arginine administration, which mitigates brain oxidative stress and enhances mitochondrial function ^30^. These findings suggest that the dysregulation of Erdr1 and Mid1 in senescence may contribute to age-related decline, highlighting their potential role as risk factors for aging.

#### 1.2 Erdr1 and Mid1 are involved in Alzheimer’s Disease pathogenesis

Alzheimer’s disease (AD) is the most common cause of dementia in older adults. It is a progressive neurodegenerative disorder characterized by abnormal processing of amyloid precursor protein (APP) and hyperphosphorylated tau proteins, leading to hallmark amyloid plaques and neurofibrillary tangles ^1^. Recent studies have implicated Mid1 in AD pathogenesis ^31–33^. Elevated Mid1 expression has been observed in AD tissues ^33^, where it enhances APP protein synthesis and upregulates BACE1, an amyloidogenic enzyme ^31,32^. Inhibiting the Mid1 protein complex has emerged as a potential therapeutic strategy to slow AD progression by reducing APP synthesis and processing, as well as lowering hyperphosphorylated tau levels ^33^. Furthermore, Erdr1 expression was found to be downregulated in mice carrying the Apolipoprotein E4 (APOE4) allele—a well-established genetic risk factor for AD—compared to mice with the non-risk APOE2 allele^34^. These findings suggest that dysregulation of Erdr1 and Mid1 may contribute to AD pathogenesis, reinforcing their potential as both risk factors and therapeutic targets.

#### 1.3 Erdr1 and Mid1 are highly involved in Parkinson’s Disease

Parkinson’s disease (PD) is a progressive neurodegenerative disorder primarily affecting movement control due to the degeneration of dopamine-producing neurons in the brain ^2^. Several studies have identified Mid1 as a potential contributor to PD pathogenesis. Specifically, two independent studies reported significantly elevated Mid1 expression in the brain tissues of PD patients, suggesting its potential role in PD progression ^35,36^. Emerging evidence also links Erdr1 to PD. Erdr1 expression was significantly downregulated in Parkin knockout mice^37^, a well-established PD mouse model characterized by increased oxidative stress and neurodegeneration. In contrast, Erdr1 levels were markedly elevated in MLKL knockout mice, which exhibit resistance to PD-like symptoms^38^. Moreover, Erdr1 is abundantly expressed in dopaminergic neurons of the A10 region, which are more resistant to degeneration, while its expression is low in the A9 region, which is more vulnerable to neurodegeneration in both PD and toxin-induced models ^39^. These findings suggest that reduced Erdr1 expression may increase susceptibility to PD, while elevated Erdr1 levels may exert a neuroprotective effect. However, conflicting evidence exists. In a 6-hydroxydopamine (6-OHDA)-induced neurotoxic mouse model of PD, Erdr1 was found to be upregulated in brain tissues^40^. This discrepancy suggests that Erdr1 may play dual roles in PD pathogenesis, potentially contributing to both neuroprotection and stress response mechanisms. Overall, these findings indicate that the dysregulation of Mid1 and Erdr1 is highly involved in PD pathogenesis.

#### 1.4 Erdr1 and Mid1 contribute to Huntington’s Disease pathogenesis

Huntington’s disease (HD) is a neurodegenerative genetic disorder caused by an expansion of CAG repeats in the HTT gene, leading to progressive nerve cell degeneration in the brain and resulting in motor, cognitive, and psychiatric symptoms ^4^. The disease occurs when the number of CAG repeats exceeds 36, while normal individuals typically carry 10 to 35 repeats ^4^. Mid1 has been shown to enhance the translation of mutant HTT mRNA in a CAG repeat-length-dependent manner ^41^. This mechanism is not unique to HD but is also observed in other disorders associated with CAG repeat expansions ^42^, suggesting that Mid1 plays a broader role in the RNA-mediated toxicity pathway ^42,43^. Emerging evidence also implicates Erdr1 in HD^44^. Huntington-null embryonic stem (ES) cell lines, which lack HTT mRNA, exhibit significantly elevated Erdr1 expression compared to wild-type counterparts ^44^. Additionally, mutant HTT ES cells exhibit altered Erdr1 levels compared to wild-type cells^44^, indicating that Erdr1 is involved in modulating both native and mutant HTT expression. Together, these findings suggest that both Mid1 and Erdr1 contribute to HD pathogenesis.

#### 1.5 Erdr1 and Mid1 are dysregulated in Amyotrophic Lateral Sclerosis

Amyotrophic lateral sclerosis (ALS) is a progressive neurodegenerative disorder that primarily affects motor neurons in the brain and spinal cord, leading to muscle weakness and loss of voluntary control ^3^. ALS is broadly classified into sporadic ALS (sALS) and familial ALS (fALS) based on its etiology. Mutations in Superoxide dismutase 1 (SOD1), a crucial antioxidant enzyme, are among the most well-characterized genetic causes of familial ALS (fALS)^45^. Interestingly, an in vitro study showed that NSC-34 cells (a widely used ALS research model) harboring mutant SOD1 exhibit reduced Erdr1 expression ^46^. Furthermore, recent studies have revealed a marked upregulation of Mid1 in the motor neurons of ALS patients^47^, as well as in ALS-sensitive motor neurons in mouse models^48^. These findings implicate Erdr1 and Mid1 in ALS pathogenesis.

#### 1.6 Erdr1 and Mid1 are strongly implicated in muscle degeneration

Muscle degenerative diseases, including myopathies and muscular dystrophies, are characterized by progressive muscle weakness and loss of muscle mass. Erdr1 and Mid1 have been shown dysregulated in mouse models exhibiting impaired muscle integrity and increased muscle atrophy^49–54^. Duchenne muscular dystrophy (DMD) is a progressive muscle degeneration disorder caused by mutations in the dystrophin gene. Elevated Erdr1 levels have been observed in dystrophin-deficient mouse models ^52^, while Mid1 dysregulation has been reported in DMD patients ^55^. Notably, Erdr1 dysregulation was observed in Auf1 knockout (KO) satellite cells, which exhibit skeletal muscle wasting and impaired regeneration following injury ^56^. Since Auf1 encodes a heterogeneous nuclear ribonucleoprotein (hnRNP) involved in pre-mRNA processing, this suggests that dysregulated Erdr1 pre-mRNA processing mediated by Auf1 may contribute to muscle degeneration pathogenesis. Together, these findings suggest that Erdr1 and Mid1 play crucial roles in muscle degenerative diseases.

#### 1.7 Erdr1 and Mid1 are strongly associated with Pulmonary Fibrosis

Pulmonary fibrosis (PF), also known as lung fibrosis, is characterized by the thickening and scarring of lung tissue, leading to a progressive decline in lung function ^6^. In the bleomycin (BLM)-induced lung fibrosis mouse model, Erdr1 expression was significantly reduced ^57^. Notably, restoring Erdr1 led to a striking reversal of established pulmonary fibrosis ^57^. Conversely, Mid1 expression was elevated in a bleomycin-induced pulmonary fibrosis mouse model and Mid1 protein levels were significantly upregulated in lung tissue of idiopathic pulmonary fibrosis (IPF) patients ^58^. Interestingly, in a radiation-induced lung fibrosis mouse model, both Erdr1 and Mid1 were upregulated in the lung tissue ^59^. Together, these findings suggest that Erdr1 and Mid1 play significant roles in the progression of pulmonary fibrosis.

#### 1.8 Erdr1 and Mid1 are critical risk factors in Rheumatoid Arthritis

Rheumatoid arthritis (RA) is a chronic inflammatory disorder characterized by persistent inflammation of the synovial joints, leading to progressive subchondral bone destruction^7^. Despite advances in therapeutic interventions, subchondral bone damage remains irreversible^7^, underscoring the urgent need for novel disease-modifying strategies. Mid1 was found to be upregulated in the synovial tissue of rheumatoid arthritis (RA) patients as well as in the collagen-induced arthritis (CIA) mouse model ^60^. Additionally, elevated Mid1 expression was observed in synovial myeloid cells and bulk knee joint tissue of the antigen-induced arthritis (AIA) model ^61^. Notably, Mid1 upregulation correlated with increased local expression of proinflammatory cytokines, including IL-1β, IL-6, and TNF, suggesting its potential involvement in the inflammatory response associated with arthritis ^62^. In contrast, in the AIA model, Erdr1 was found to be downregulated in synovial myeloid cells and bulk knee joint tissue ^61^. In the CIA model, recombinant Erdr1 (Erdr1-177) demonstrated therapeutic effects by inhibiting IL-18 production and suppressing synovial fibroblast migration ^63^. Furthermore, both Erdr1 and Mid1 were markedly suppressed in synovial myeloid cells of Fas-deficient mice, which exhibited resistance to joint destruction following AIA induction compared to wild-type mice ^62^. Together, these findings suggest that Erdr1 and Mid1 play significant roles in RA pathogenesis and may serve as potential therapeutic targets.

#### 1.9 Erdr1 and Mid1 are key factors driving diabetes and its complications

Diabetes is marked by chronic hyperglycemia due to defects in insulin secretion, action, or both, leading to damage to insulin-producing β cells ^8^. Erdr1 and Mid1 have been implicated in promoting insulin production in β cells under hyperglycemic conditions ^64^. However, their overactivation in response to hyperglycemia may contribute to β cell destruction, suggesting a potential role in diabetes progression.

Beyond their involvement in insulin regulation, Erdr1 and Mid1 were also involved in diabetes-related complications. Erdr1 expression was significantly downregulated in the kidneys of db/db diabetic mice and involved in diabetic kidney disease (DKD) ^65^. Notably, effective DKD therapies, including angiotensin receptor blockers (ARBs) and sodium-glucose cotransporter 2 inhibitors (SGLT2i), restored Erdr1 expression, thereby protecting the kidneys from injury in a mouse model^65^. Similarly, Erdr1 expression was dramatically restored in pancreatic islets and peripheral lymph nodes (PLNs) following beneficial treatment (Tyrosine protein-kinase 2 (TYK2) inhibitor treatment) in a non-obese diabetic (NOD) mouse model ^66^. These findings suggest a protective role of Erdr1 in diabetes.

However, Erdr1 expression, along with Mid1, was reported to be elevated in the brains of NOD mice ^67^. Moreover, Mid1 was implicated in renal injury associated with DKD, as its expression was found to be elevated in DKD patients ^68^. In a db/db diabetic mouse model, DKD symptoms were alleviated following Mid1 knockdown via Mid1 shRNA adenovirus injection ^68^. *In vitro,* Mid1 knockdown inhibited high glucose-induced inflammation and diabetic fibrosis in the HK-2 cell line^68^. These findings suggest that Erdr1 and Mid1 are risk factors for diabetes and its complications.

Collectively, these findings suggest that Erdr1 and Mid1 play crucial roles in the pathogenesis of diabetes and diabetes-related complications, highlighting their potential as promising therapeutic targets for diabetes management.

#### 1.10 Conclusion: Erdr1 and Mid1 are shared risk factors for aging and degenerative diseases

These studies demonstrate that upregulated Mid1 is a common risk factor for aging and degenerative diseases. However, some studies suggest that upregulated Erdr1 serves as a risk factor, others indicate that downregulated Erdr1 contributes to disease pathogenesis. Overall, we conclude that Erdr1 and Mid1 act as shared risk factors for aging and degenerative diseases (**Figure 1**). We propose that they contribute to these conditions by regulating oxidative stress. Our study aims to investigate the role of Erdr1 and Mid1 in modulating oxidative stress and cellular damage.

**Figure 1.**
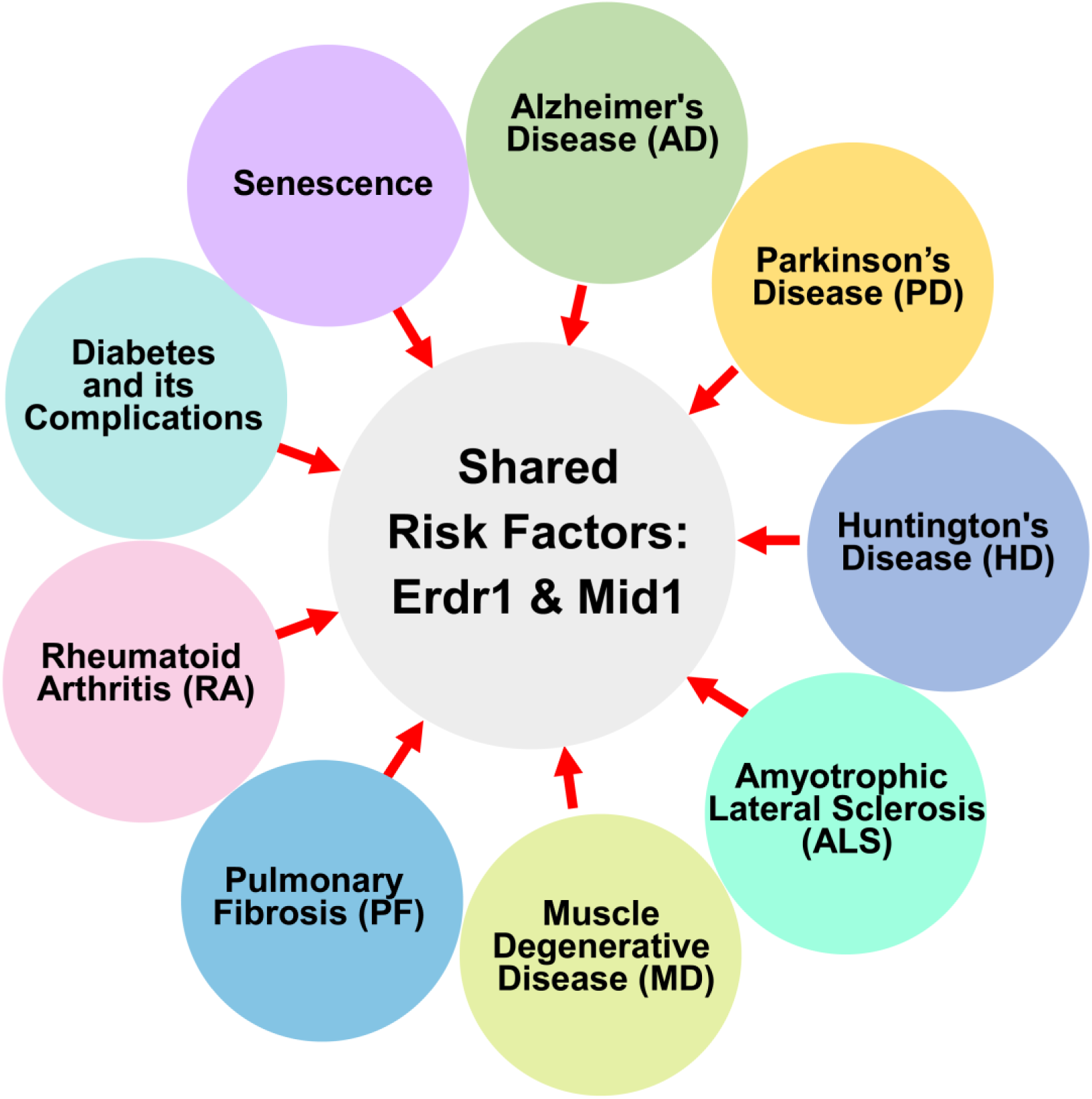
Erdr1 and Mid1 are shared risk factors for multiple degenerative diseases.

### 2. Erdr1 exhibits alternative splicing upon oxidative stress

#### 2.1 Erdr1 predominantly exists in three isoforms

In mice, Erdr1 is located on the sex chromosomes, neighboring Mid1, and is encoded by distinct gene loci on the X chromosome (Gm21887) and the Y chromosome (Gm47283). The UniProtKB database identifies mainly three isoforms of Erdr1: Erdr1-145 (Primary Accession No. Q6PED5), Erdr1-177 (Primary Accession No. O88727), and Erdr1-209 (Primary Accession No. Q811B0). Our data reveal that the purified proteins of these three Erdr1 isoforms exist as monomers and exhibit distinct molecular weights of approximately 22 kDa, 26 kDa, and 30 kDa, respectively (**Figure 2A**). When overexpressed in 293T cells, these isoforms exhibited similar monomer types and sizes (data not shown).

**Figure 2.**
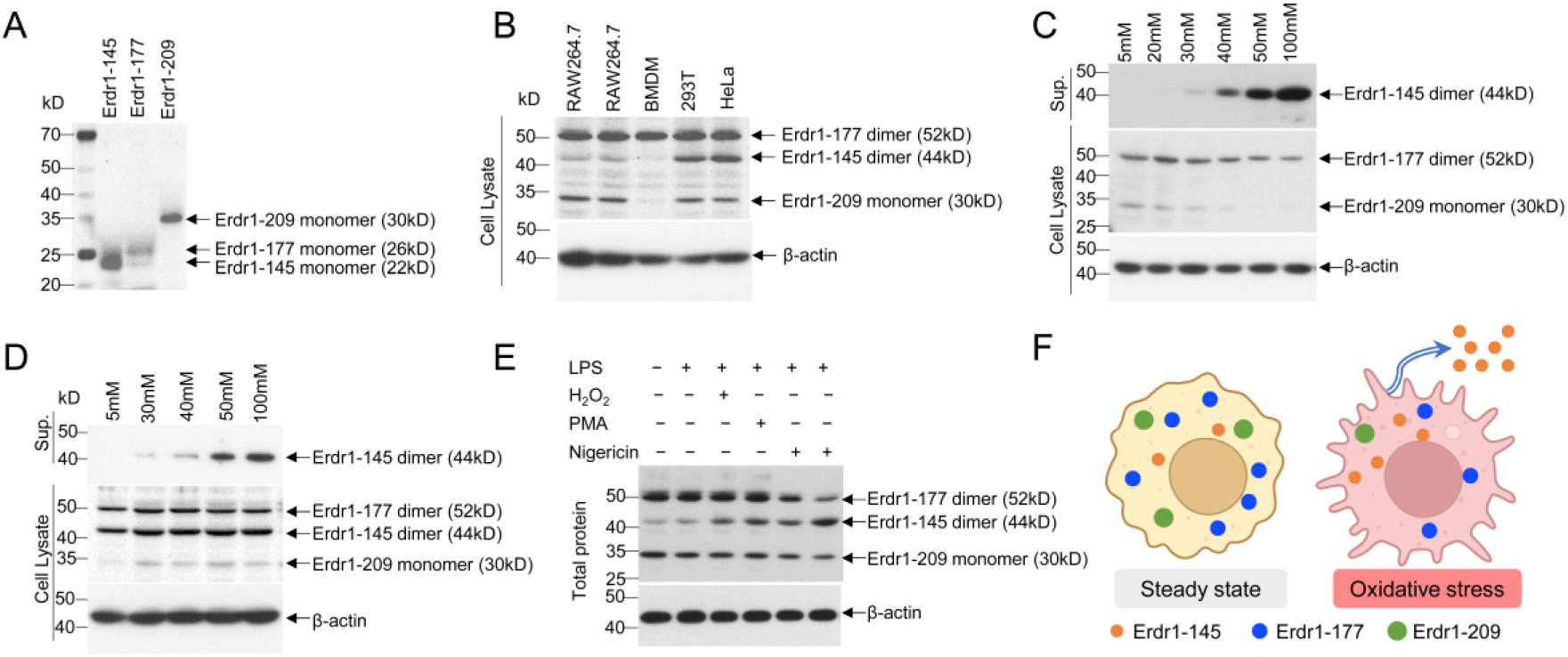
Erdr1 functions as a sensor for cellular oxidative Stress through alternative splicing. A. Monomer sizes of the three Erdr1 isoforms: Erdr1-145 (∼22 kDa) and Erdr1-177 (∼26 kDa), Erdr1-209 (∼ 30kDa). B. Western blot analysis of Erdr1 expression in indicated cell lines. β-actin was detected as an internal control. C. Western blot analysis of Erdr1 expression in cell lysates and supernatants of RAW 264.7 cells treated with indicated glucose concentrations for 24 hours. D. Western blot analysis of Erdr1 expression in cell lysates and supernatants of HeLa cells treated with indicated glucose concentrations for 24 hours. E. Western blot analysis of Erdr1 expression in RAW 264.7 cells treated with the indicated oxidative stress inducers for 1–4 hours. Cells were pre-primed either without LPS (Sample 1) or with 500 ng/mL LPS for 3 hours (Samples 2–6). The indicated samples contained both cells and supernatant, collectively referred to as total protein. Sample 3: 500 μM H2O2; Sample 4: 10 μM PMA; Sample 5:10 uM Nigericin; Sample 6: 20 uM Nigericin. F. A model of Erdr1 response to oxidative stress. In response to oxidative stress, Erdr1 undergoes alternative splicing, leading to the downregulation of the Erdr1-177 and Erdr1-209 isoforms while promoting the secretion of the Erdr1-145 isoform. Results are representative of three independent experiments.

#### 2.2 Erdr1 exhibits distinct basal expression patterns in different cell lines

We analyzed basal Erdr1 expression across various cell lines. In RAW 264.7 cells, Erdr1-177 dimers (about 52 kD) are the predominant isoforms, while Erdr1-209 monomers (about 30 kD) and Erdr1-145 dimers (about 44 kD) are expressed at lower levels (**Figure 2B**). In contrast, both Erdr1-177 dimers and Erdr1-145 dimers emerge as dominant isoforms in HeLa and 293T cells (**Figure 2B**). These findings suggest that Erdr1 exhibits unique basal expression patterns in different cell lines.

#### 2.3 Erdr1 undergoes alternative Splicing in response to oxidative stress

We next investigated whether Erdr1 functions as a sensor for oxidative stress, with a particular focus on high glucose, a well-established inducer of cellular oxidative stress. Previously, we observed a reduction in Erdr1-177 dimer levels following LPS treatment in RAW 264.7 cell ^24^. Here, we further treated RAW 264.7 cells with high glucose concentrations (20–100 mM) for 24 hours, following a 3-hour LPS priming step. Our results indicate that Erdr1-177 dimers, along with Erdr1-209, were significantly downregulated in a glucose dose-dependent manner. In contrast, Erdr1-145 dimers exhibited a robust increase and were actively secreted (**Figure 2C**). We also observed a marked increase in the release of Erdr1-145 dimers in response to high glucose in HeLa cells (**Figure 2D**) and 293T cells (data not shown). Additionally, other oxidative stress inducers, such as H2O2, PMA, and Nigericin, also triggered Erdr1-145 secretion in RAW 264.7 cells (**Figure 2E**). Collectively, these findings indicate that Erdr1 undergoes alternative splicing in response to oxidative stress, with the upregulation and secretion of Erdr1-145 emerging as a common phenomenon in different cell lines (**Figure 2F**).

### 3. Erdr1 isoforms distinctly regulate oxidative stress

The structures of the three isoforms of Erdr1 were predicted using AlphaFold2 (**Figure 3A–B**). Erdr1-177 and Erdr1-209 exhibited similar structural conformations (**Figure 3A–B**). Erdr1-177 has been demonstrated to play an anti-inflammatory role ^24^, and our preliminary data suggest that Erdr1-209 also exerts anti-inflammatory effects. In contrast to Erdr1-177 and Erdr1-209, Erdr1-145 has a distinct structural configuration (**Figure 3A–B**) and functions as a pro-inflammatory isoform ^24^. We then investigate the role of these Erdr1 isoforms in modulating oxidative stress by performing protein supplementation experiments. Our findings reveal that Erdr1-145 primes ROS production in RAW 264.7 cells and markedly enhances high glucose-induced ROS generation, whereas Erdr1-177 and Erdr1-209 exert opposing inhibitory effects (**Figure 4C**). Given that ROS is a well-established trigger of inflammatory responses and cellular damage, we further examined the impact of Erdr1 isoforms on high glucose-induced IL-1β secretion and LDH release. Our results show that Erdr1-145 enhances high glucose-induced IL-1β secretion (**Figure 4D**) and LDH release (**Figure 4E**). In contrast, Erdr1-177 and Erdr1-209 suppress these processes (**Figure 4D–E**). Collectively, these findings demonstrate that Erdr1 isoforms differentially regulate oxidative stress and cellular damage. Erdr1-177 and Erdr1-209 act as antioxidants, while Erdr1-145 functions as a pro-oxidant.

**Figure 3.**
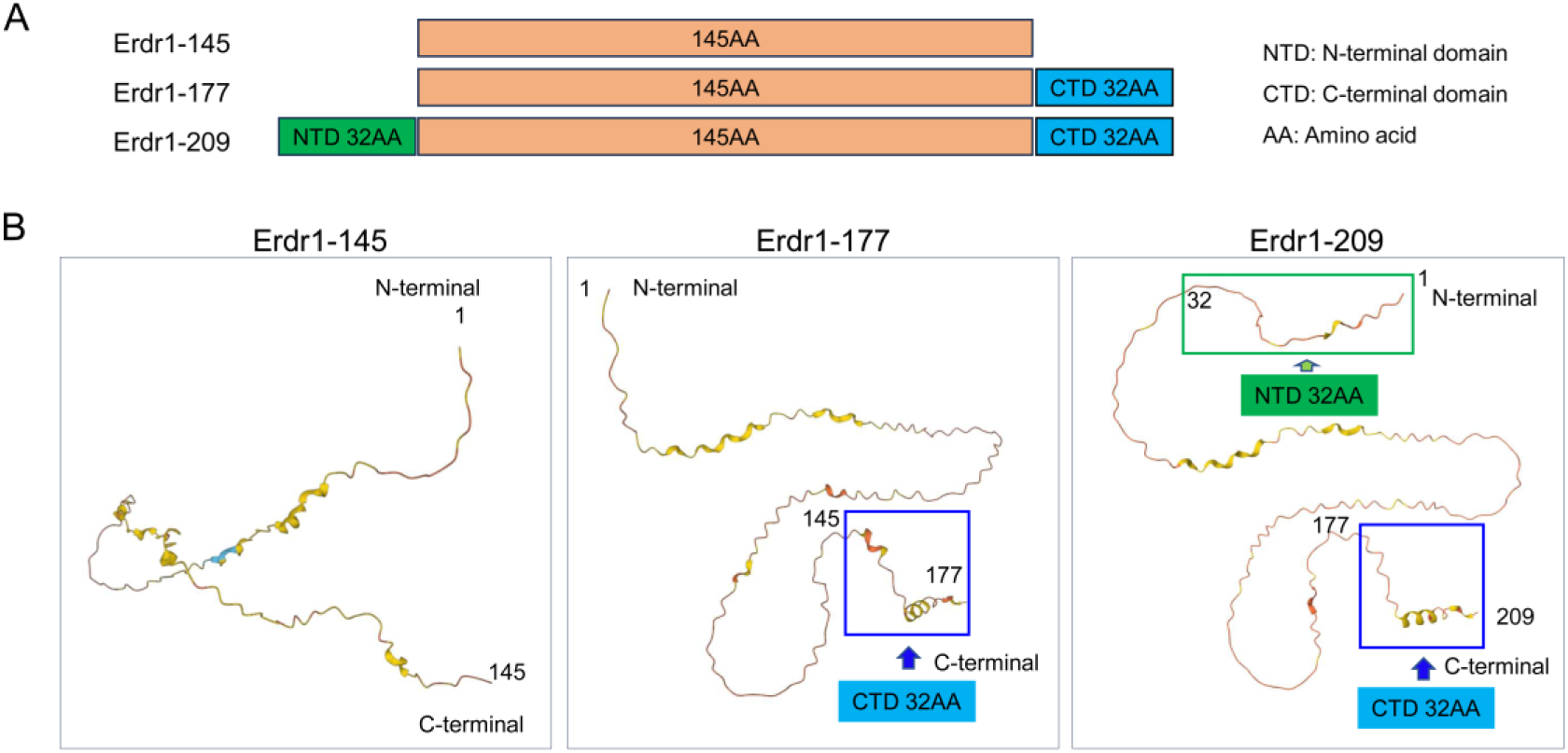
Erdr1 isoforms exhibit distinct structural characteristics. A. Structural differences of the three Erdr1 alternative splice isoforms. B. The predicted structure of three isoforms of Erdr1 by using alphafold2.

**Figure 4.**
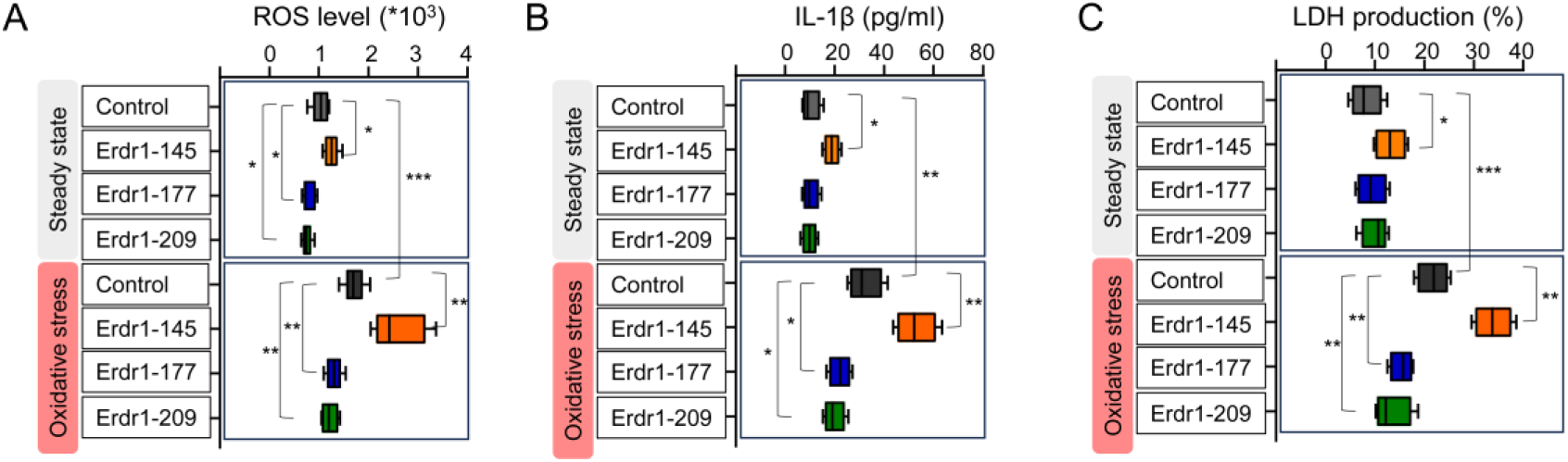
Erdr1 isoforms differentially regulate high glucose-induced oxidative stress. RAW 264.7 cells were subjected to oxidative stress by treatment with 50 mM glucose for 24 hours. Cells were additionally treated with or without 100 ng/mL of the indicated Erdr1 isoforms. Cells cultured in a normal glucose medium served as the steady-state control. A. Detection ROS production (n=6). B. ELISA for detecting IL-1β production (n=4). C. LDH production (n=4). Box plots were generated with whiskers extending from the minimum to the maximum values to represent the full range of the data. Statistical significance was determined using Student’s t-test (*p < 0.05, **p < 0.01, ***p < 0.001). Results are representative of three independent experiments.

### 4. Erdr1 isoforms modulate oxidative stress by regulating Mid1 activation

We have previously demonstrated that Mid1 serves as a downstream effector of Erdr1 in the pro-inflammatory pathway ^24^. Here, we propose that Erdr1 isoforms modulate oxidative stress by regulating Mid1 activation, which is closely linked to oxidative stress activation ^69^.

#### 4.1 Mid1 is a sensor and trigger of cellular oxidative stress

To determine whether Mid1 is a sensor of oxidative stress, we examined its activation under high glucose conditions in RAW 264.7 cells. Immunofluorescence analysis revealed that Mid1 was significantly induced by high glucose treatment (**Figure 5A–B**, panels 1-2). This finding is consistent with a previous report demonstrating elevated Mid1 expression in kidney cell lines exposed to high glucose ^68^. We further investigated the role of Mid1 in high glucose-induced oxidative stress. Our findings demonstrate that Mid1 knockout significantly reduces high glucose-induced ROS production and LDH release, whereas Mid1 overexpression enhances both processes (**Figure 5C–D**). These indicate that Mid1 serves as a sensor and trigger of cellular oxidative stress.

**Figure 5.**
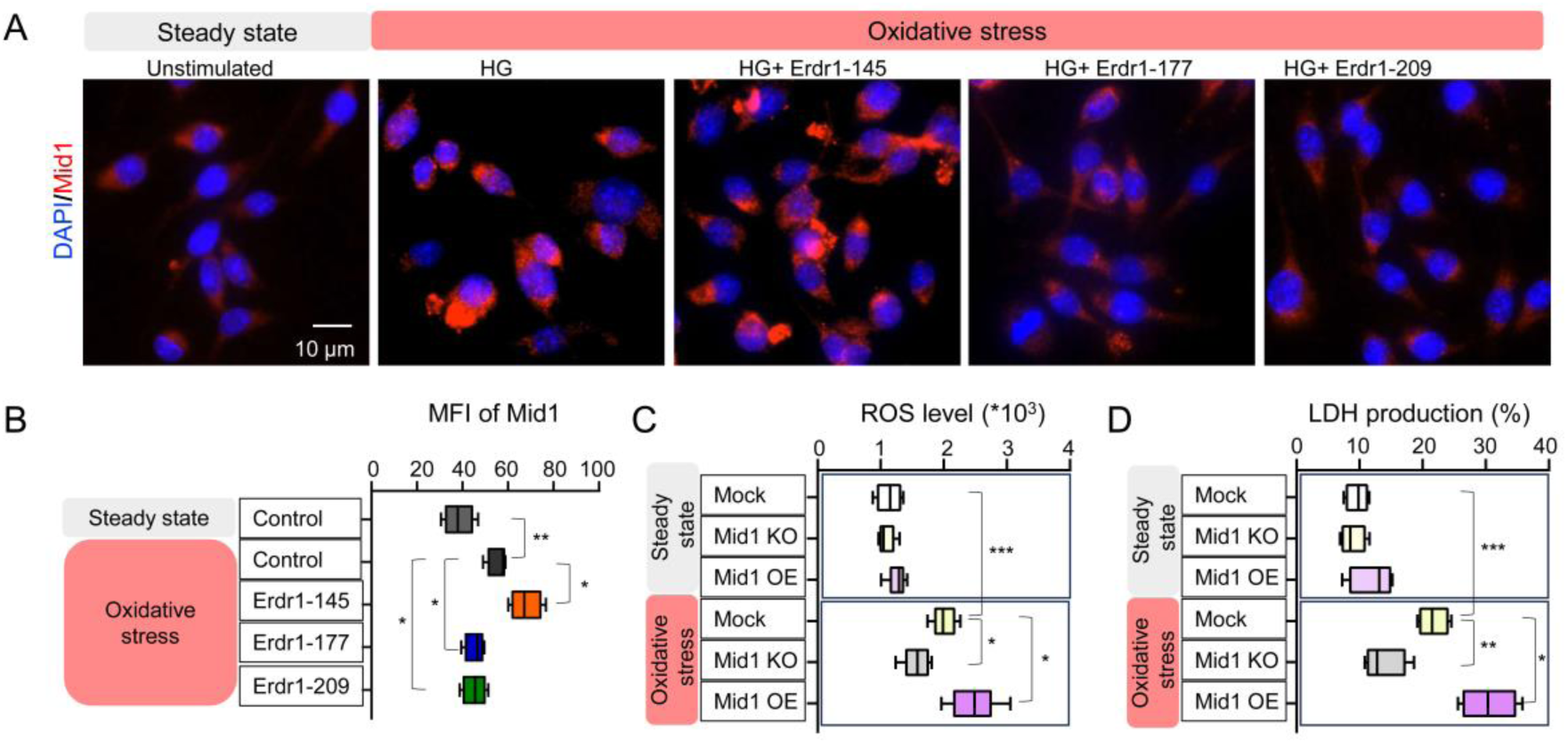
Erdr1 isoforms modulate oxidative stress by regulating Mid1 activation. A. Representative immunofluorescence images showing Mid1 expression in unstimulated (non-induced) RAW 264.7 cells and high glucose (HG)-induced RAW 264.7 cells (panels 2–5) treated with or without 100 ng/mL of the indicated Erdr1 isoforms. B. Mean fluorescence intensity (MFI) of Mid1 as quantified from (A) using ImageJ (n = 4). C. Measurement of ROS production in control, Mid1 knockout, and Mid1-overexpressing RAW 264.7 cells, with or without high glucose treatment. D. Measurement of LDH release in control, Mid1 knockout, and Mid1-overexpressing RAW 264.7 cells, with or without high glucose treatment. (A–D) Oxidative stress was induced in RAW 264.7 cells by treatment with 50 mM glucose for 24 hours. Box plots were generated with whiskers extending from the minimum to the maximum values to represent the full range of the data. Statistical significance was determined using Student’s t-test (*p < 0.05, **p < 0.01, ***p < 0.001). Results are representative of three independent experiments.

#### 4.2. Erdr1 isoforms distinctly regulate Mid1 activation

We further explored the roles of Erdr1 isoforms in regulating Mid1 activation under oxidative stress conditions. Our results demonstrate that Erdr1-145 enhances Mid1 activation in response to high glucose, whereas Erdr1-177 and Erdr1-209 significantly suppress Mid1 activation (**Figure 5A–B**). These findings establish Erdr1 as an upstream regulator of Mid1 in oxidative stress responses and highlight the distinct roles of Erdr1 isoforms in Mid1 activation. Specifically, Erdr1-177 and Erdr1-209 mitigate oxidative stress by inhibiting Mid1 activation, thereby preventing oxidative stress-induced cellular damage. In contrast, Erdr1-145 promotes oxidative stress and contributes to cellular damage by activating Mid1 (**Figure 7**). The Erdr1-Mid1 oxidative stress axis provides a potential mechanistic explanation for their roles as shared risk factors in aging and degenerative diseases.

## Discussion

### Erdr1 and Mid1 collaboratively engage in multiple biological processes

Erdr1 and Mid1 are highly interconnected and co-implicated in a broad spectrum of biological processes, including neurodegenerative diseases, neurodevelopmental disorders, brain injury, inflammation, asthma, longevity, telomere protection, hematopoietic capacity, T-cell immune regulation, cellular migration and so on (**Supplemental Table 1**). Their shared involvement in these biological processes suggests that they may contribute to a wide range of both physiological and pathological conditions.

### Erdr1 and Mid1 are highly associated with redox-related factors

In this study, we identified Erdr1 and Mid1 as shared risk factors for aging and degenerative diseases and demonstrated that their involvement in oxidative stress regulation underlies this association. Erdr1 has been reported to be strongly associated with redox systems (**Figure 6**; **Supplemental Table 2**). On the one hand, Erdr1 is linked to pro-oxidant systems, such as ROS production ^70,71^ and NADPH oxidases (e.g., Nox2, Nox4) which regulate ROS production ^52,70,72^. Additionally, Erdr1 expression is upregulated in response to H₂O₂ exposure, a well-known inducer of oxidative stress ^73^. On the other hand, Erdr1 also links to key antioxidant systems, including Superoxide Dismutase 1 (SOD1) ^46^, Vitamin B12 ^74^, glutathione system (e.g., GPX and GSH) ^23,75,76^, and thioredoxin system ^75,77^. Importantly, Erdr1 is highly interconnected with the Nrf2 signaling pathway ^75,78^, a master regulator of the antioxidant response ^79^. Similarly, Mid1 has been shown to prime oxidative stress and exacerbate liver injury ^69^. Our study further reveals that Mid1 acts as a trigger for high glucose-induced oxidative stress and cellular damage and demonstrates that Erdr1 isoforms differentially regulate oxidative stress by modulating downstream Mid1 activation. This provides a molecular basis for the Erdr1-Mid1 axis in driving oxidative damage and cellular degeneration (**Figure 7**).

**Figure 6.**
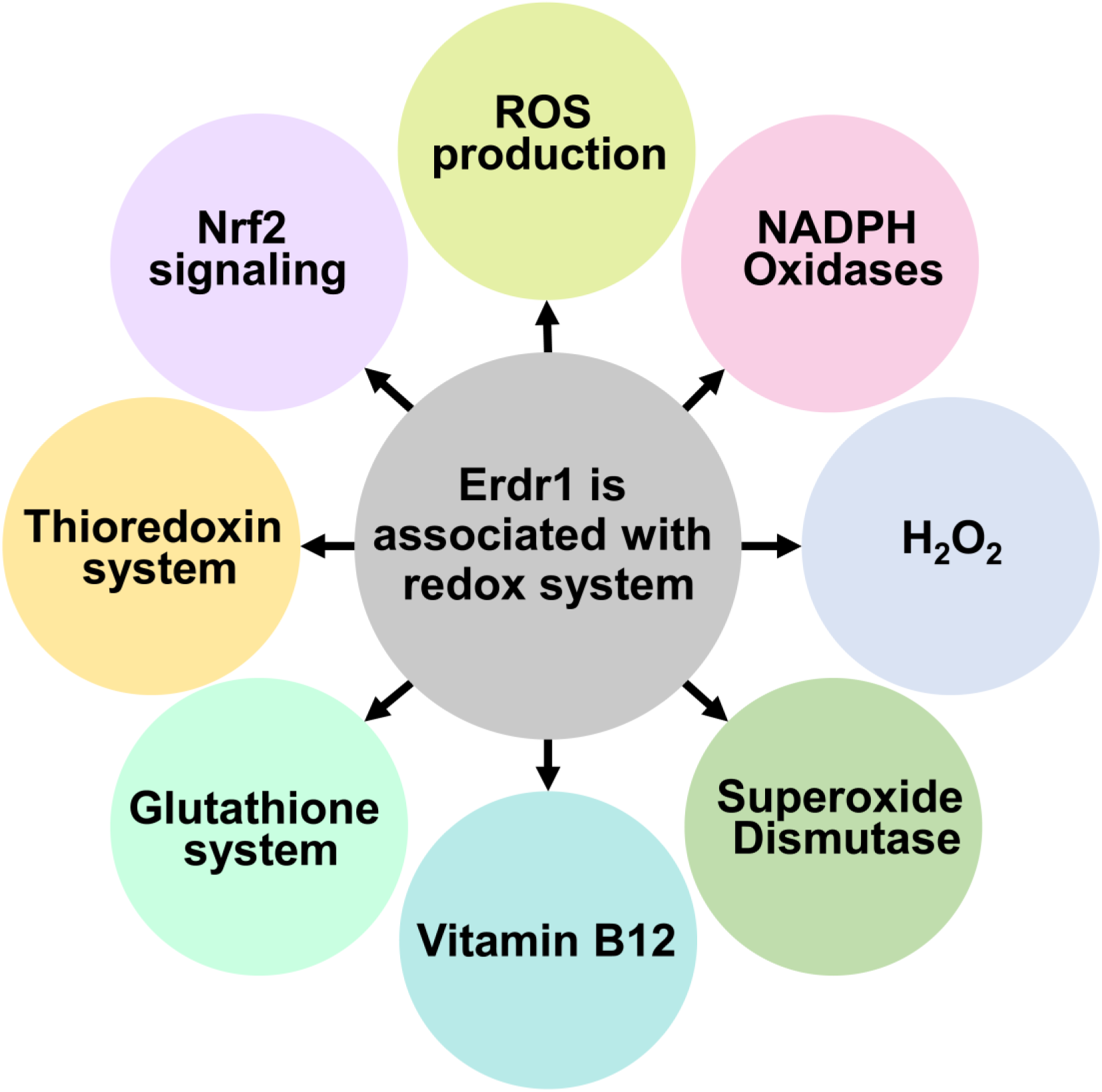
Erdr1 is associated with redox-related factors.

**Figure 7.**
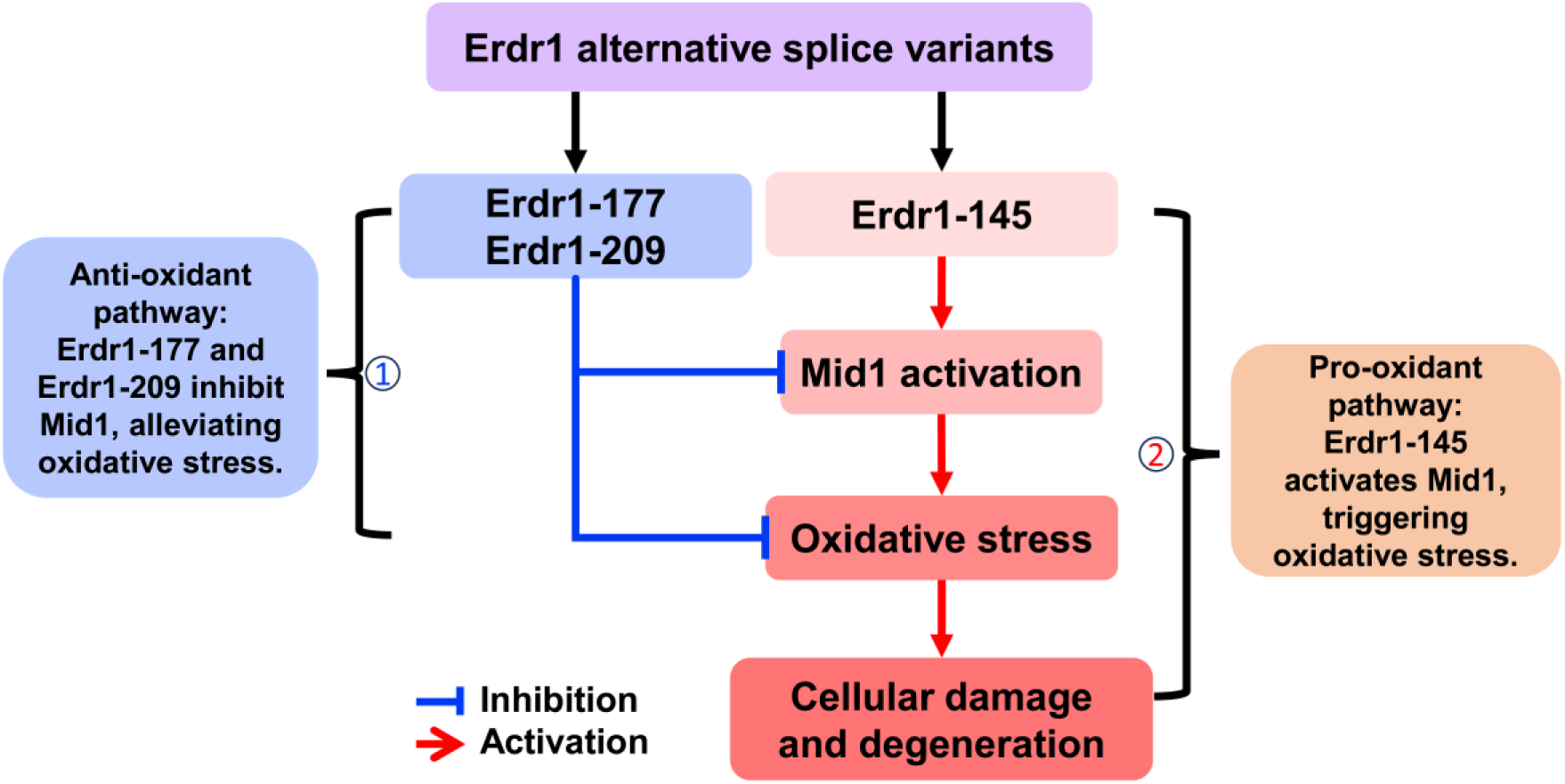
Work model of Erdr1-Mid1 axis in oxidative stress and cellular damage. The three Erdr1 alternative spliced isoforms display distinct roles in oxidative stress and cell damage. 1) Erdr1-177 and Erdr1-209 play protective roles by inhibiting Mid1 activation and mitigating oxidative stress. 2) Erdr1-145 activates Mid1, triggering oxidative stress, leading to cellular damage and degeneration.

### Erdr1 exhibits alternative splicing upon oxidative stress exposure

Previous studies have reported substantial variability in Erdr1 RNA transcripts, particularly at the 5’ end ^80,81^. Many of these transcripts lack complete open reading frames (ORFs), suggesting that Erdr1 RNA undergoes incomplete processing until translation is required ^81^. This property facilitates a rapid translational response to cellular stress and supports cellular adaptation. Our data further reveal that Erdr1 undergoes alternative splicing upon oxidative stress, altering the balance of its three isoforms (**Figure 8**). Erdr1-177 and Erdr1-209, which have antioxidant properties, are downregulated in response to oxidative stress. Erdr1-145, a pro-oxidant isoform that interacts with Mid1, is upregulated and secreted under oxidative stress conditions. This alternative splicing mechanism increases Erdr1’s functional complexity and reinforces its diverse roles in oxidative stress regulation.

**Figure 8.**
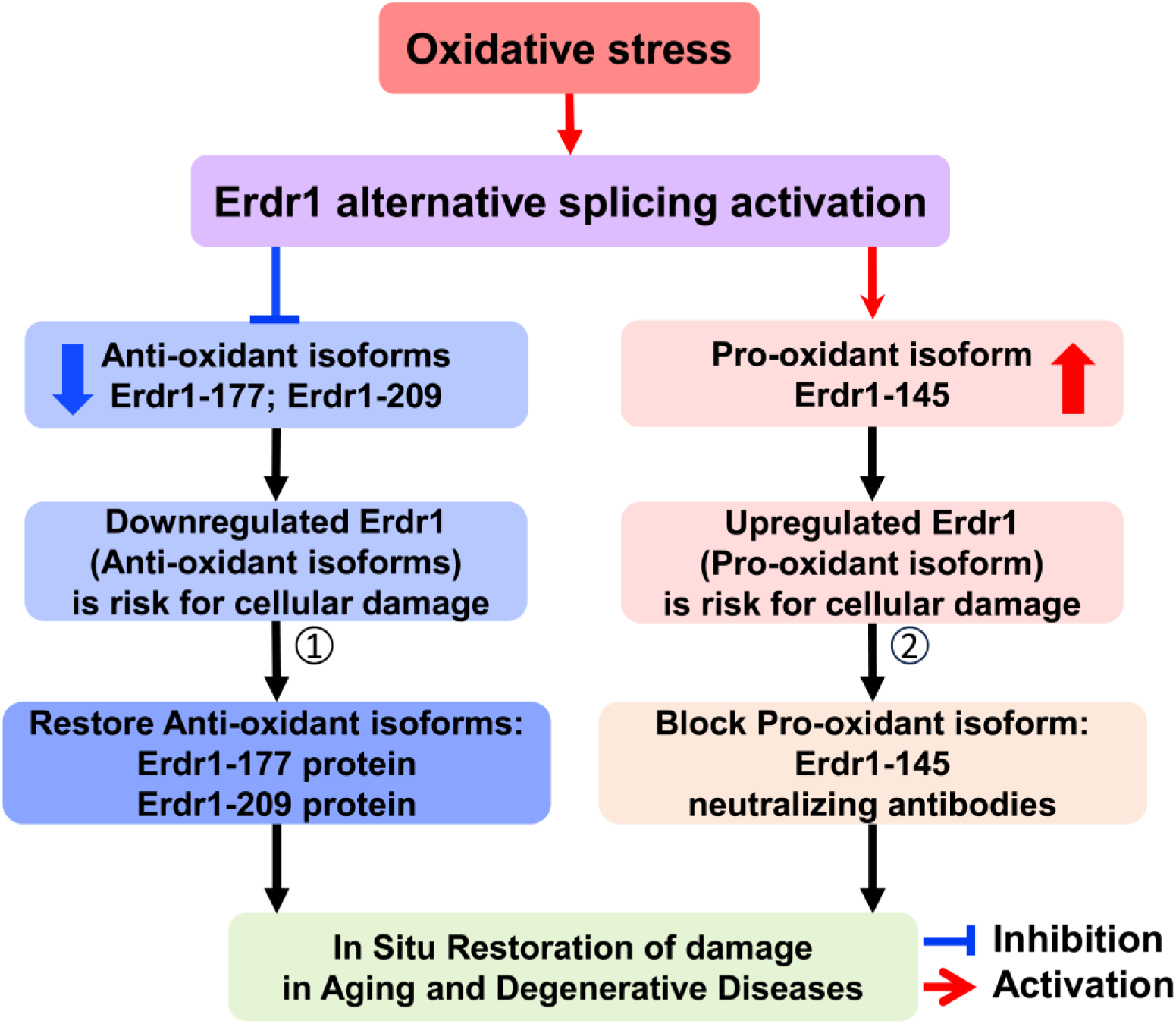
Strategies for in situ restoration of damage in aging and degenerative diseases by modulating Erdr1. Erdr1 mediates cellular responses to oxidative stress through alternative splicing, resulting in the suppression of antioxidant isoforms Erdr1-177 and Erdr1-209, while promoting the production of the pro-oxidant isoform Erdr1-145. Dysregulation of Erdr1 contributes to cell damage and degeneration (**as shown in Figure 6**). This model explains how both downregulated Erdr1 (Erdr1-177 and Erdr1-209) and upregulated Erdr1 (Erdr1-145) are risk factors for aging and degenerative diseases (**Figure 1**, **Table 1**). Here, we propose two strategies for in situ restoration of damage in aging and degenerative diseases by modulating Erdr1. 1). Restoring antioxidant isoform levels by supplementing with Erdr1-177 or Erdr1-209 proteins. 2). Inhibiting the pro-oxidant isoform by designing and utilizing Erdr1-145 neutralizing antibodies.

### Key finding: First identification of Erdr1-145

A key finding of this study is the discovery of Erdr1-145 as an oxidative stress-induced secreted isoform (**Figure 2F**). This represents the first identification of this novel Erdr1 isoform, highlighting its pro-oxidant role as an emerging cytokine. Erdr1-145 contributes to pro-inflammatory responses, oxidative stress, and cellular damage, which collectively drive cellular senescence. Senescent cells exhibit a distinctive secretory profile, known as the Senescence-Associated Secretory Phenotype (SASP) ^82^. Our findings suggest that Erdr1-145 likely serves as a key contributor to SASP, further implicating it in aging and degenerative processes.

### In situ restoration for aging and degenerative diseases by modulating Erdr1

Dysregulation of Erdr1 isoforms in response to oxidative stress—characterized by a decrease in antioxidant isoforms (Erdr1-177 and Erdr1-209) or an increase in the pro-oxidant isoform (Erdr1-145)—contributes to oxidative stress and cellular damage (**Figure 7–8**). This finding highlights potential therapeutic strategies to modulate Erdr1 to prevent oxidative stress-induced damage in aging and degenerative diseases. We propose two potential approaches.

**1. Restoring the expression of antioxidant isoforms.** This can be achieved by supplementing with Erdr1-177 or Erdr1-209 proteins to counteract oxidative stress.
**2. Inhibiting pro-oxidant Isoforms.** Developing Erdr1-145 neutralizing antibodies to mitigate oxidative damage.

These strategies suggest the potential for in situ regeneration and functional restoration of damaged cells, tissues, and organs in aging and degenerative diseases (**Figure 8**).

### Limitations of current research and future directions

Mid1 has consistently been found to be upregulated at the RNA level in all degenerative mouse models we have documented, with some findings also confirmed at the protein level ^33,58,68^. Elevated levels of Mid1 have likewise been observed in patient samples of certain degenerative diseases ^33,35,36,58,60,68^. Moreover, Mid1 inhibition has shown protective effects against degeneration in both in vitro and in vivo models ^60,68^, supporting its strong potential as a therapeutic target. However, the association between Erdr1 and aging and degenerative diseases remains primarily based on RNA sequencing data from mouse models. Several critical gaps remain, requiring further investigation to validate these findings.

**1. Validation at the protein level**: It is crucial to determine whether RNA-level changes in Erdr1 are reflected at the protein level in cells and animal models. Given Erdr1’s alternative splicing, it is essential to monitor Erdr1 isoforms dysregulation in degenerative tissues.
**2. Functional analysis of Erdr1 isoforms in mouse models:** Future studies should focus on overexpressing distinct Erdr1 isoforms rather than simply knocking out the Erdr1 gene. This approach will clarify the unique roles of each isoform in aging and degenerative diseases.
**3. Establishing a direct link between Erdr1 and human degenerative conditions:** Although Erdr1 mRNA and protein sequences are completely conserved between humans and mice^80,81^, and we observed Erdr1 expression in human cell lines (**Figure 1B**), no data are currently available regarding Erdr1 expression in human diseases due to the absence of a mapped human Erdr1 gene ^24^. Mapping the human Erdr1 gene and confirming its expression in human patients is necessary to establish clinical relevance.

## Conclusion

In conclusion, this study demonstrates that Erdr1 and Mid1 function as shared risk factors for aging and various degenerative diseases. Our findings highlight that Erdr1-145 induces oxidative stress and cellular damage by activating Mid1. This provides a molecular mechanistic explanation for the shared role of Erdr1 and Mid1 in aging and degenerative diseases. Furthermore, we propose therapeutic strategies to mitigate cellular damage by regulating Erdr1 levels, suggesting a straightforward and effective approach for in situ repair of damage associated with aging and degenerative diseases.

## Authors’ Contributions

Y.W. designed and performed the experiments, prepared the figures, interpreted the data, and wrote the manuscript. D.P. contributed to experimental design, data interpretation, and manuscript revision. N.L. contributed to experimental execution, data interpretation, and manuscript revision.

## Acknowledgments

Many thanks to K.W.P. for his truly unparalleled and invaluable contribution to this groundbreaking idea. His remarkable curiosity and unique perspective have been a profound source of inspiration.

## Abbreviations

AD: Alzheimer’s disease
ALS: Amyotrophic Lateral Sclerosis
AIA: Antigen-induced arthritis
BLM: Bleomycin
CIA: Collagen-induced arthritis
DKD: Diabetic kidney disease
DMD: Duchenne muscular dystrophy
Erdr1: Erythroid Differentiation Regulator 1
HD: Huntington’s disease
IPF: Idiopathic pulmonary fibrosis
LDH: Lactate Dehydrogenase
MD: Muscular Dystrophy
Mid1: Midline 1
NOD: Non-obese diabetic
PD: Parkinson’s disease
PF: Pulmonary Fibrosis
RA: Rheumatoid Arthritis
ROS: reactive oxygen species
SAMP: Senescence-Accelerated Mouse Prone
SASP: Senescence-Associated Secretory Phenotype
SOD1: Superoxide Dismutase 1

