## Supplemental Table 1 and 2, Figure 1 and 2 for "Exploring Erdr1-Mid1 Axis: Shared Risk Factors and Molecular Mechanisms in Aging and Degenerative Diseases"

**Supplemental Table 1.**
**Erdr1 and Mid1 collaboratively engage in multiple biological processes**

| Processes involved | Erdr1 and Mid1 expression | Article title | Publish Year | Reference |
|---|---|---|---|---|
| Neurodegenerative disease | Erdr1 and Mid1 were downregulated in the brains of PPT1 KO mice, a mouse model of infantile neuronal ceroid lipofuscinosis. | Gene expression profiling in a mouse model of infantile neuronal ceroid lipofuscinosis reveals upregulation of immediate early genes and mediators of the inflammatory response | 2007 | PMID: 18021406 |
| | Erdr1 and Mid1 were downregulated in Chrnb2 KO mice. | Mouse Mutants for the Nicotinic Acetylcholine Receptor ß2 Subunit Display Changes in Cell Adhesion and Neurodegeneration Response Genes | 2011 | PMID: 21547082 |
| | Erdr1 and Mid1 were downregulated in mice carrying AD-risked humanized APOE4 compared to those with APOE2. | Humanized APOE genotypes influence lifespan independently of tau aggregation in the P301S mouse model of tauopathy | 2023 | PMID: 37337279 |
| Neurodevelopmental disorder | Erdr1 and Mid1 were retained in 39, XY*O mice, which is a genetic mouse model of neurodevelopmental disorder. | Biological mechanisms associated with increased perseveration and hyperactivity in a genetic mouse model of neurodevelopmental disorder | 2012 | PMID: 23276394 |
| | Erdr1 was significantly reduced in the brain of 39, XY*O mice. | Altered brain gene expression but not steroid biochemistry in a genetic mouse model of neurodevelopmental disorder | 2014 | PMID: 24602487 |
| | Erdr1 and Mid1 were upregulated in macrophages of FMRP deficiency mice which exhibits neurodevelopmental disorder. | The Fragile X Syndrome-Related Protein, FMRP, Modulates Innate Immune Gene Expression in Macrophages | 2022 | the FASEB journal<br><br>P51OD011104 |

| | | | | |
|---|---|---|---|---|
| | Erdr1 and Mid1 were upregulated in the brain of a mouse model of prenatal PCB exposure. | Placenta and fetal brain share a neurodevelopmental disorder DNA methylation profile in a mouse model of prenatal PCB exposure | 2022 | PMID: 35235788 |
| Brain Injury | Erdr1 and Mid1 were downregulated in the CNS of SARM KO mice which exhibited reduced CNS injury and cytokine production upon virus infection. | SARM Is Required for Neuronal Injury and Cytokine Production in Response to Central Nervous System Viral Infection | 2013 | PMID: 23749635 |
| | Erdr1 and Mid1 were downregulated after effective injury treatment. | Repetitive Mild Traumatic Brain Injury and Transcription Factor Modulation | 2020 | PMID: 32292111 |
| Rheumatoid Arthritis | Erdr1 was downregulated, and Mid1 was upregulated in myeloid cells of mice after AIA (antigen-induced arthritis) induction. | Mid1 is a novel mediator of subchondral bone resorption in antigen-induced arthritis | 2019 | 4th annual meeting of the European Calcified Tissue Society |
| | Erdr1 and Mid1 were both downregulated in myeloid cells of Fas KO mice which are resistant to joint destruction. | Myeloid-specific molecular mediators of subchondral bone damage in antigen-induced arthritis | 2021 | Poster code: R-01-06-115 |
| Inflammation | Erdr1 and Mid1 were upregulated in inflammatory response. | The miR-23a~27a~24-2 microRNA Cluster Promotes Inflammatory Polarization of Macrophages | 2021 | PMID: 33328213 |
| | Erdr1 was downregulated, and Mid1 was upregulated in M1 macrophages. | Erdr1 Drives Macrophage Programming via Dynamic Interplay with YAP1 and Mid1 | 2024 | PMID: 38392560 |
| | Erdr1 and Mid1 were downregulated in the microglia of mice after vitamin B12 treatment which alleviates neuroinflammation and injury. | Functional regulation of microglia by vitamin B12 alleviates ischemic stroke-induced neuroinflammation in mice. | 2024 | PMID: 38715940 |

| | | | | |
|---|---|---|---|---|
| Asthma | Erdr1 and Mid1 were both downregulated in the lung tissue of IgM KO mice which has reduced airway hyperresponsiveness. | Immunoglobulin M regulates airway hyperresponsiveness independent of T helper 2 allergic inflammation | 2023 | Elife (https://doi.org/10.7554/eLife.90531.1) |
| Lung fibrosis | Erdr1 and Mid1 were upregulated by radiation-induced lung disease. | Gene expression profiles among murine strains segregate with distinct differences in the progression of radiation-induced lung disease | 2017 | PMID: 28130353 |
| Longevity, telomere protection and hematopoietic capacity | Erdr1 and Mid1 were upregulated in LBR-deficient mice which exhibited reduced lymphocyte longevity and homeostatic proliferation. | Reduced Lymphocyte Longevity and Homeostatic Proliferation in Lamin B Receptor-Deficient Mice Results in Profound and Progressive Lymphopenia | 2012 | PMID: 22105998 |
| | Erdr1 and Mid1 were downregulated after TERRA depletion which exhibited increased telomerase activity. | TERRA RNA Antagonizes ATRX and Protects Telomeres | 2017 | PMID: 28666128 |
| | Erdr1 and Mid1 were downregulated in the bone marrow of Neil3 KO mice which displayed decreased hematopoietic capacity and reduced telomere length. | NEIL3-deficient bone marrow displays decreased hematopoietic capacity and reduced telomere length | 2022 | PMID: 35079641 |
| T cell Immune regulation | Erdr1 and Mid1 were both up-regulated in YAP1 KO Tregs. | YAP Is Essential for Treg-Mediated Suppression of Antitumor Immunity | 2018 | PMID: 29907586 |

| | | | | |
|---|---|---|---|---|
| Insulin production and diabetes | Erdr1 and Mid1 were upregulated in the islets of SLC30A8 R138X mice, which exhibited increased insulin production in response to hyperglycemia. | Mice harboring the human SLC30A8 R138X loss-of-function mutation have increased insulin secretory capacity | 2018 | PMID: 30038024 |
| | Erdr1 and Mid1 were upregulated in the brains of NOD mice, a well-established animal model for type 1 diabetes. | NOD mouse dorsal root ganglia display morphological and gene expression defects before and during autoimmune diabetes development | 2023 | PMID: 37334284 |
| Cellular migration | Erdr1 and Mid1 expression were dramatically downregulated in CCR5 deficiency pulmonary mesenchymal cells. | CC-Chemokine Receptor 5 on Pulmonary Mesenchymal Cells Promotes Experimental Metastasis via the Induction of Erythroid Differentiation Regulator 1 | 2014 | PMID: 24197118 |
| | Erdr1 was upregulated and Mid1 was downregulated in IL-10 deficiency B-1 lymphocytes. | The axis IL-10/claudin-10 is implicated in the modulation of aggressiveness of melanoma cells by B-1 lymphocytes | 2017 | PMID: 29145406 |
| Development | Erdr1 and Mid1 were upregulated in placental tissue of Csf2 KO mice, which exhibited altered placental development. | Csf2 Null Mutation Alters Placental Gene Expression and Trophoblast Glycogen Cell and Giant Cell Abundance in Mice | 2009 | PMID: 19228596 |
| Obesity | SNPs and indels in Erdr1 and Mid1 are associated with obesity | Whole genome sequencing of mouse lines divergently selected for fatness (FLI) and leanness (FHI) revealed several genetic variants as candidates for novel obesity genes | 2024 | PMID: 38483771 |
| Spermatogenesis Male Infertility | Erdr1 and Mid1 were upregulated in the testis of SCCx43KO mice, which exhibited infertile phenotype. | Sertoli-cell-specific knockout of connexin 43 leads to multiple alterations in testicular gene expression in prepubertal mice | 2012 | PMID: 22699423 |

**Supplemental Table 2. Erdr1 is strongly associated with Redox-related factors (related to Figure 6)**

| Redox-related factors | Erdr1 expression | Article title | Publish Year | Reference |
|---|---|---|---|---|
| ROS | Erdr1 was downregulated in the liver of Hvcn1-deficient mice, where Hvcn1 plays a crucial role in regulating ROS production. | Regulation of hepatic oxidative stress by voltage-gated proton channels (Hv1/VSOP) in Kupffer cells and its potential relationship with glucose metabolism. | 2020 | PMID: 33040408 |
| | Erdr1 and Hvcn1 were the most upregulated genes in the islets of SLC30A8 R138X mice, which showed enhanced insulin production in response to hyperglycemia. | Mice harboring the human SLC30A8 R138X loss-of-function mutation have increased insulin secretory capacity | 2018 | PMID: 30038024 |
| | Erdr1 was the most downregulated gene in the hearts of miR-100 overexpression mice, which exhibited reduced ROS production. | Cardiomyocyte-specific miR-100 overexpression preserves heart function under pressure overload in mice and diminishes fatty acid uptake as well as ROS production by direct suppression of Nox4 and CD36 | 2021 | PMID: 34605573 |

| | | | | |
|---|---|---|---|---|
| NADPH oxidases | Erdr1, along with Nox4 and Lox, was upregulated in the left ventricles of mdx mice (dystrophin-deficient mice). | Dystrophin-deficient cardiomyopathy in mouse: expression of Nox4 and Lox are associated with fibrosis and altered functional parameters in the heart | 2008 | PMID: 18440230 |
| | Erdr1 was downregulated in Nox2 KO mice compared to wild-type mice after ischemic preconditioning followed by ischemia-reperfusion | Novel role of NADPH oxidase in ischemic myocardium: a study with Nox2 knockout mice. | 2012 | PMID: 22038056 |
| | Erdr1 and Nox4 were the most downregulated genes in the hearts of miR-100 overexpression mice, which showed reduced ROS production. | Cardiomyocyte-specific miR-100 overexpression preserves heart function under pressure overload in mice and diminishes fatty acid uptake as well as ROS production by direct suppression of Nox4 and CD36. | 2021 | PMID: 34605573 |
| $H_2O_2$ | Erdr1 was upregulated and ROS production was enhanced in cardiomyocytes treated with $H_2O_2$. | Suppression of lncRNA Gm47283 attenuates myocardial infarction via miR-706/ Ptgs2/ferroptosis axis. | 2022 | PMID: 35485136 |
| SOD1 | Erdr1 expression was downregulated in NSC-34 cells, an ALS model cell line harboring mutant SOD1. | Mutant SOD1 alters the motor neuronal transcriptome: implications for familial ALS. | 2005 | PMID: 15872021 |
| Vitamin B12 | Erdr1 was downregulated in microglia of mice after vitamin B12 treatment which alleviates neuroinflammation and injury. | Functional regulation of microglia by vitamin B12 alleviates ischemic stroke-induced neuroinflammation in mice. | 2024 | PMID: 38715940 |

| | | | | |
|---|---|---|---|---|
| Glutathione system | Overexpression of Erdr1 in T cells suppressed the expression of GPX7 and GPX4. | Microbiota promotes systemic T-cell survival through suppression of an apoptotic factor | 2017 | PMID: 28487480 |
| | Erdr1 was dramatically downregulated, whereas GPX2 was significantly elevated in TrxR1/Gsr-null livers. | TrxR1, Gsr, and oxidative stress determine hepatocellular carcinoma malignancy | 2019 | PMID: 31097586 |
| | Knockdown of Erdr1 in Neuro2a cells promotes GSH synthesis. | Erythroid Differentiation Regulator 1 as a Regulator of Neuronal GSH Synthesis. | 2024 | PMID: 39061840 |
| Thioredoxin system | Erdr1 was significantly upregulated in Txnrd1 deficiency T cells. | The thioredoxin-1 system is essential for fueling DNA synthesis during T-cell metabolic reprogramming and proliferation. | 2018 | PMID: 29749372 |
| | Erdr1 was dramatically downregulated in TrxR1/Gsr-null livers. | TrxR1, Gsr, and oxidative stress determine hepatocellular carcinoma malignancy | 2019 | PMID: 31097586 |
| Nrf2 signaling | Erdr1, along with the Nrf2 target genes Nqo1 and Srxn1, were significantly upregulated in Txnrd1-deficient T cells. | The thioredoxin-1 system is essential for fueling DNA synthesis during T-cell metabolic reprogramming and proliferation. | 2018 | PMID: 29749372 |
| | Erdr1 was dramatically downregulated, whereas Nrf2-response genes (Aox1, Gsta4, Gstm3, Srxn1, Nqo1, Cbr3, etc.) were significantly upregulated in TrxR1/Gsr-null livers. | TrxR1, Gsr, and oxidative stress determine hepatocellular carcinoma malignancy | 2019 | PMID: 31097586 |
| | Erdr1 was dramatically upregulated in the lungs of Nrf2 deficiency mice. | Nrf2 Regulates Granuloma Formation and Macrophage Activation during Mycobacterium avium Infection via Mediating Nramp1 and HO-1 Expressions. | 2021 | PMID: 33563837 |

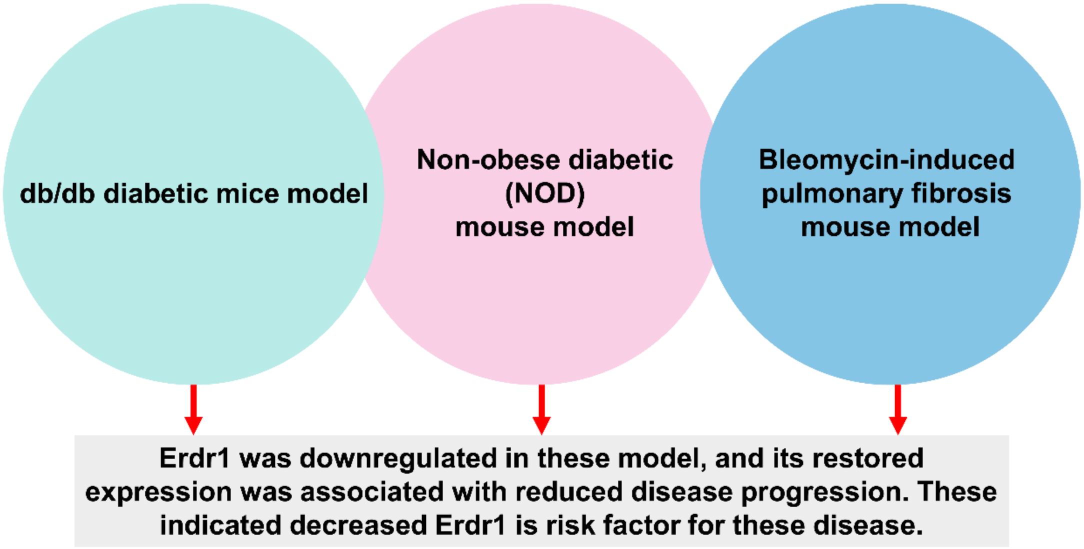


**Supplemental Figure 1. Downregulated Erdr1 is indicated as a common risk factor for these diseases.**

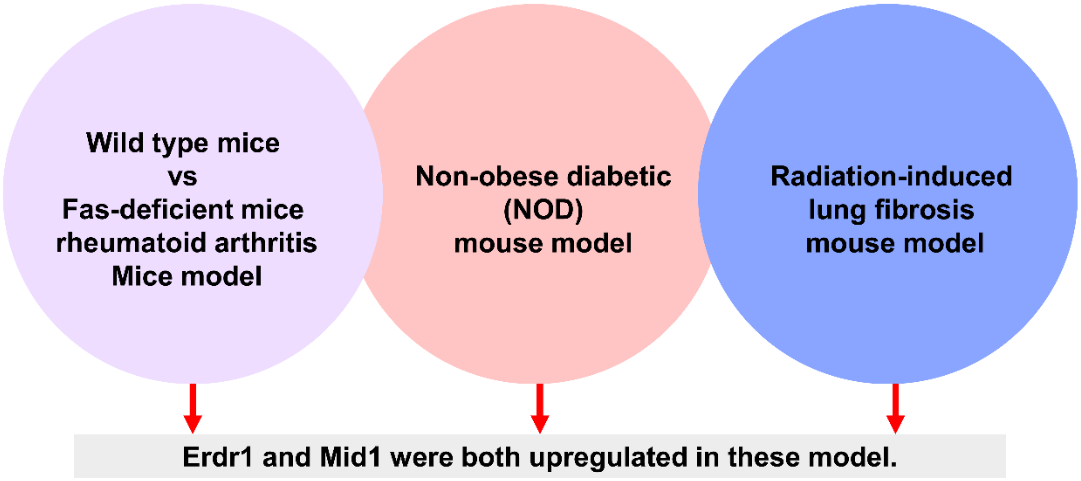


**Supplemental Figure 2. Both Erdr1 and Mid1 are upregulated in these disease models.**
